# Modulation of Arabidopsis growth by volatile organic compounds from a root-derived bacterial community

**DOI:** 10.1101/2022.04.12.488003

**Authors:** Gözde Merve Türksoy, Réjane Carron, Anna Koprivova, Stanislav Kopriva, Kathrin Wippel, Tonni Grube Andersen

**Affiliations:** Max Planck Institute for Plant Breeding Research, Carl-von-Linne-Weg 10, 50829, Cologne, Germany; Cluster of Excellence on Plant Sciences, University of Cologne, 50674 Cologne, Germany; Institute for Plant Sciences, University of Cologne, 50674 Cologne, Germany; Germany

## Abstract

Plant roots are surrounded by fluctuating biotic and abiotic factors. The living component – the microbiota – is actively shaped by the plant and plays an important role in overall plant health. While it has been shown that specialized metabolites exuded from the plant are involved in shaping host interactions with the microbiota, it is unclear how underground volatile organic compounds (VOCs) influence this communication. This is especially true for root-associated bacteria which are known to release VOCs that can influence plant growth. Using a simplified synthetic bacterial community (SynCom) representing the phylogenetic diversity of bacteria in the root microbiome, we set out to characterize plant growth and defense metabolites when subjected to bacterial VOCs (bVOCs). Moreover, by profiling the SynCom community composition after co-cultivation with the plant, we explored how members of the community influenced each other in our growth setup. Our findings reveal that plant growth promotion can occur via VOCs from a bacterial SynCom, but that the plant response differs for individual community members. In addition, we find that bVOCs are able to repress chemical defense responses in the plant, possibly to facilitate colonization. By removing key species from the SynCom, we find that complex bacteria-bacteria interactions are likely to underlie this phenomenon, and that bVOC-induced modulation of plant responses in the rhizosphere may be an emergent property of bacterial communities rather than depending on individual species.

## Introduction

In nature, roots of healthy plants associate with a diverse set of bacteria, fungi, oomycetes and nematodes, collectively termed the root microbiota (Bai et al., 2015; Bulgarelli et al., 2013). These microbial communities – in particular bacteria – have been shown to be crucial for host health, protection against pathogens, and nutrient acquisition from the surrounding soil (Castrillo et al., 2017; Harbort et al., 2020; Trapet et al., 2016; J. Zhang et al., 2019). Dependent on their needs, plants are able to actively modulate the microbiome through induction of plant defenses (Carrión et al., 2019) or exudation of metabolites (Bais et al., 2004; Tyc, et al., 2017; Zhalnina et al., 2018), which, in certain cases can promote plant growth (Durán et al., 2018). Thus, the assembly and structure of microbial communities are dynamic and rely on active communication with the host plant (Bulgarelli et al., 2012; Lundberg et al., 2012; Wippel et al., 2021). Arabidopsis and other species of the Brassicales order employ a specialized set of sulfur-containing amino-acid-derived metabolites as a remarkably efficient defense system (Künstler et al., 2020). In Arabidopsis they are mainly derived from methionine, which gives rise to the aliphatic glucosinolates, as well as from tryptophan for indolic glucosinolates and camalexin, which partake in shaping the root microbiome (Bednarek, 2012; Koprivova et al., 2019; Siebers et al., 2018).

While such specialized plant metabolites help to shape the bacterial microbiome (Jacoby et al., 2021; Wolinska et al., 2021), the role of underground metabolites from the bacterial communities in plant-microbiota interaction is less clear. In particular, bacteria release Volatile Organic Compounds (bVOCs) that are characterized by a low molecular mass (< 300 Da), low boiling point and high vapor pressure allowing interaction and signaling in short- and long-distance via soil, air and water (Schulz & Dickschat, 2007; Weisskopf et al., 2021). bVOC production and composition is dependent on the growth substrate and environment (Blom et al., 2011; Fincheira & Quiroz, 2018). Yet, a number of strains have been reported to modulate plant growth via volatiles, leading to increases in biomass and alteration of the root system architecture (RSA) (Groenhagen et al., 2013; C.-M. Ryu et al., 2003; Syed Ab Rahman et al., 2018). Moreover, bVOCs have been found to induce plant defenses or directly inhibit pathogen growth, resulting in reduced disease severity (Chinchilla et al., 2019; Lee et al., 2012; Piechulla et al., 2017; C. M. Ryu et al., 2004). These growth modulations have been linked to specific volatile compounds and their interference with the host phytohormone balance (Bailly et al., 2014; Tahir et al., 2017; Tyagi et al., 2019; H. Zhang et al., 2007). Most bVOCs-related studies have been conducted with single bacterial strains or with low-diversity bacterial communities with a focus on community richness effects (Raza et al., 2020). Thus, we lack a framework for understanding bVOCs-mediated modulations in a close-to-nature bacterial community context (Weisskopf et al., 2021). Quantifiable synthetic communities (SynComs) as representatives of the phylogenetic bacterial diversity in natural soils provide a powerful tool to further analyze the role of bVOCs for plant-microbe communication, but also in microbial community assembly and maintenance.

Here we describe the bVOCs-mediated A. thaliana growth modulation in the context of a previously established 16-member SynCom, which recapitulates growth-promoting effects found under root-colonizing conditions in soil-based systems (Durán et al., 2018; Wippel et al., 2021). We assessed modulation capacity of the SynCom and of the 16 individual members on plant growth as well as bacterial community structure. Our work identified individual strains that affected plant growth differentially in direct bacterial contact (DBC), in presence of spatially separated bacteria (no bacterial contact, NBC), or combined treatments. Combined, our findings indicate that bVOCs may modulate plant growth independently and cumulatively, in a strain-dependent manner. Thus, plant growth modulation may be an emergent property of the bVOC bouquet released by the community, which is likely to play an important role in bacteria-bacteria interaction in a community context.

## Material and Methods

### Plant material and growth conditions

Arabidopsis thaliana wild type ecotype Columbia-0 (Col-0) was used for all experiments. Seeds were surface-sterilized for 5 min with 70% EtOH, followed by brief wash of 100% EtOH. Dried sterilized seeds were sowed on ½ strength Murashige & Skoog medium (1/2 MS) without sucrose (2.45 g/L MS salts including vitamins and MES buffer (Duchefa), 8 g/L Bacto Agar (BD Biosciences), pH 5.8) square plates (Greiner, 120×120 mm). Plates were sealed with Micropore tape, placed in the dark at 4°C for 2 days for stratification, then transferred to a growth chamber vertically (16 h light/ 8 h dark cycle, 21/19°C) for germination for 5 days.

### Bacterial growth conditions

The 16 strains from SynCom At-SC3 (Wippel et al., 2021, Root1221 replaced by Root29, Root1485 replaced by Root418) and WCS417, a beneficial Pseudomonas strain (Pieterse et al.,2021) were used in this study. Bacterial strains were cultivated on solid medium containing 15 g/L tryptic soy broth (TSB; Sigma-Aldrich) and 15 g/L Bacto Agar and incubated at 25°C. Information on the individual bacterial strains can be found at At-RSPHERE (http://www.at-sphere.com/) (Bai et al., 2015). Prior to the start of the experiment, liquid cultures were started from single bacterial colonies in 4 mL liquid TSB and incubated shaking overnight at 25°C.

### Co-cultivation of plants and bacteria

For the non-bacterial contact (NBC) and direct bacterial contact (DBC) plate setup, 1/2 MS medium containing 10 g/L Bacto Agar was poured into square petri dishes (Greiner,120×120mm), with a small petri dish lid (Greiner, 35×10 mm) positioned at the middle of the bottom (Supplementary Figure 1A). The small dish was filled with 4 mL TSB agar medium and allowed to solidify. Five-day-old Col-0 seedlings were placed approximately 1 cm below the top of the square plate system, and for NBC or combined treatments 100 μl of bacterial culture (OD600=1, in 0.9 % sodium chloride) were added to the small petri dish lid. For mock controls and DBC experiments, 0.9 % NaCl was inoculated on the TSB plate. For the DBC conditions, we followed a previously established protocol (Ma et al., 2021). Briefly, strains were adjusted to OD600 = 0.2 using 0.9% NaCl, and 150 μl of bacterial culture were added to 50 ml 40°C warm MS agar medium to yield a final density of OD=0.0005, and poured into a square petri dish. Plates were stored at room temperature overnight before seedling transfer. For mock treatment, 150 μl 0.9% NaCl were added into 50 ml 40°C warm MS agar medium. For dilution experiments, the total bacterial culture was adjusted to OD600=0.25, 0.5 or 1.0. To prepare the SynComs, 50 μl each of the 16 bacterial strains (OD600=1) were combined and vortexed briefly. All plates were sealed with Micropore tape after inoculation. Pictures were taken after 10 days of exposure to bVOCs (15 days old seedlings). The shoot fresh weight was measured at harvesting, and primary root length as well as number of lateral roots were quantified from the photographs using manual measurements in Image J (Fiji) (Schindelin et al., 2012).

### Metabolite analysis

Camalexin was extracted from ca. 20 mg shoot tissue in 100 μL of dimethylsulfoxide (DMSO) for 20 min with shaking and after centrifugation, 20 μL were injected into a Thermo Scientific Dionex UltiMate 3000 HPLC system with Waters Spherisorb ODS-2 column (250 mm x 4.6 mm, 5 μm). The samples were resolved using a gradient of 0.01% (v/v) formic acid (solvent A) and a solvent mixture of 98% (v/v) acetonitrile 2% (v/v) water and 0.01% (v/v) formic acid (solvent B). The gradient program was as follows: 97% A for 5 min; 90% A in 5 min; 40% A in 8 min; 20% A in 2 min; 0% A in 20 min and kept at 0% A for 10 min; 100% A in 2.5 min and kept 100% A for 3.5 min; 97% A in 2 min and kept 97%A for 2 min. Camalexin was detected at an excitation at 318 nm and emission at 368 nm by fluorescence (FLD sensitivity set to 3) exactly as described in Koprivova et al., 2019. Glucosinolate content was determined using reverse phase HPLC with UV detection as described by Dietzen et al., 2020

### Community profiling

DNA from bacteria was isolated via alkaline lysis. A streak of bacteria was resuspended in 200 μL of 0.9% NaCl, and 12 μL of the sample were added to 20 μL of Buffer I (25 mM NaOH, 0.2 mM EDTA(Na), pH 12), mixed by pipetting, and incubated at 94 °C for 30 min. 20 μL of Buffer II (40 mM Tris-HCl, pH 7.46) was added to neutralize pH. Bacterial communities were profiled by amplicon sequencing of the variable v5-v7 regions of the bacterial 16S rRNA gene. Library preparation for Illumina MiSeq sequencing was performed as described previously (Durán et al., 2018). In all experiments, multiplexing of samples was performed by double-indexing (barcoded forward and reverse oligonucleotides for 16S rRNA gene amplification). Amplicon sequencing data was demultiplexed according to their barcode sequence using the QIIME pipeline (Caporaso et al., 2010) Quality-filtered merged paired-end reads were then aligned to a reference set of sequences extracted from the whole-genome assemblies of every strain included in a given SynCom, using Rbec (v1.0.0). A count table was generated that was employed for downstream analyses of diversity in R (v4.0.3) with the R package vegan (v2.5–6). Finally, we visualized amplicon data from all experimental systems using the ggplot2 R package (v3.3.0) (Wickham, 2016). To quantitatively compare the relative abundance of individual strains in the drop-out SynComs vs. the 16-member SynCom, the sequencing reads of the omitted strain were removed from the count table of the full SynCom samples prior to calculation of relative abundances.

### Statistical analysis

Pairwise comparisons were performed using a two-tailed Student’s t-test in Microsoft Excel (treatment mean vs. control mean). GraphPad version 8.02 was used to conduct one-way ANOVA with a Tukey’s HSD (honestly-significant-difference) post-hoc pairwise T-test. Letters and asterisks (*) indicate statistically significant difference of means (p < 0.01). Graphs were generated using GraphPad version 8.02 or R. For boxplots, the centre bar depicts the median and the lower and upper box limits depict the 25th and 75th percentile, respectively, whiskers represent minima and maxima, closed dots depict individual samples.

## Results

### VOCs from a beneficial bacterial SynCom have opposing effects on plant root and shoot growth

To initiate our analyses, we employed a plate-in-plate growth setup that allows physical separation of plants and bacteria but with shared headspace. In this setup, bacteria are confined to grow on an open small petri dish embedded at the bottom of a bigger square plate which is sealed from the outside with Micropore tape (Supplementary Figure 1A). To study bVOC effects on Arabidopsis seedlings, we chose a previously established 16-member SynCom consisting of strains originally isolated from roots of Arabidopsis grown in a natural soil and covering a broad taxonomic range (Bai et al., 2015; Wippel et al., 2021) (Supplementary Figure 1B). By cultivating plants alone (mock), with bacteria mixed in the plant growth medium, or bacteria in the separate compartment, we assessed to what extend this SynCom can modulate plant growth by direct bacterial contact (DBC) or non-bacterial contact (NBC). In line with previous findings (Durán et al., 2018; Wippel et al., 2021), in the DBC configuration, the SynCom gave rise to a significant increase in the total fresh weight of plants compared to mock-treated control (p<0.01) (Figure 1A, D). Intriguingly, in the NBC and the combined configuration, we saw a similar, significant (P<0.01), increase in fresh weight (Figure. 1A, D), suggesting that bVOCs may be involved in the growth promotion mediated by this SynCom. However, both NBC and combined treatments led to a significant decrease in root length and a concurrent increase in lateral root occurrence (P<0.01), whereas no significant changes were observed for either of these measurements in the DBC-treated roots (Figure 1B, C and D). Thus, it is likely that the bVOC bouquet arising from a community of root-associated bacteria affects the above- and belowground tissues of the host plant differently.

**Figure 1.**
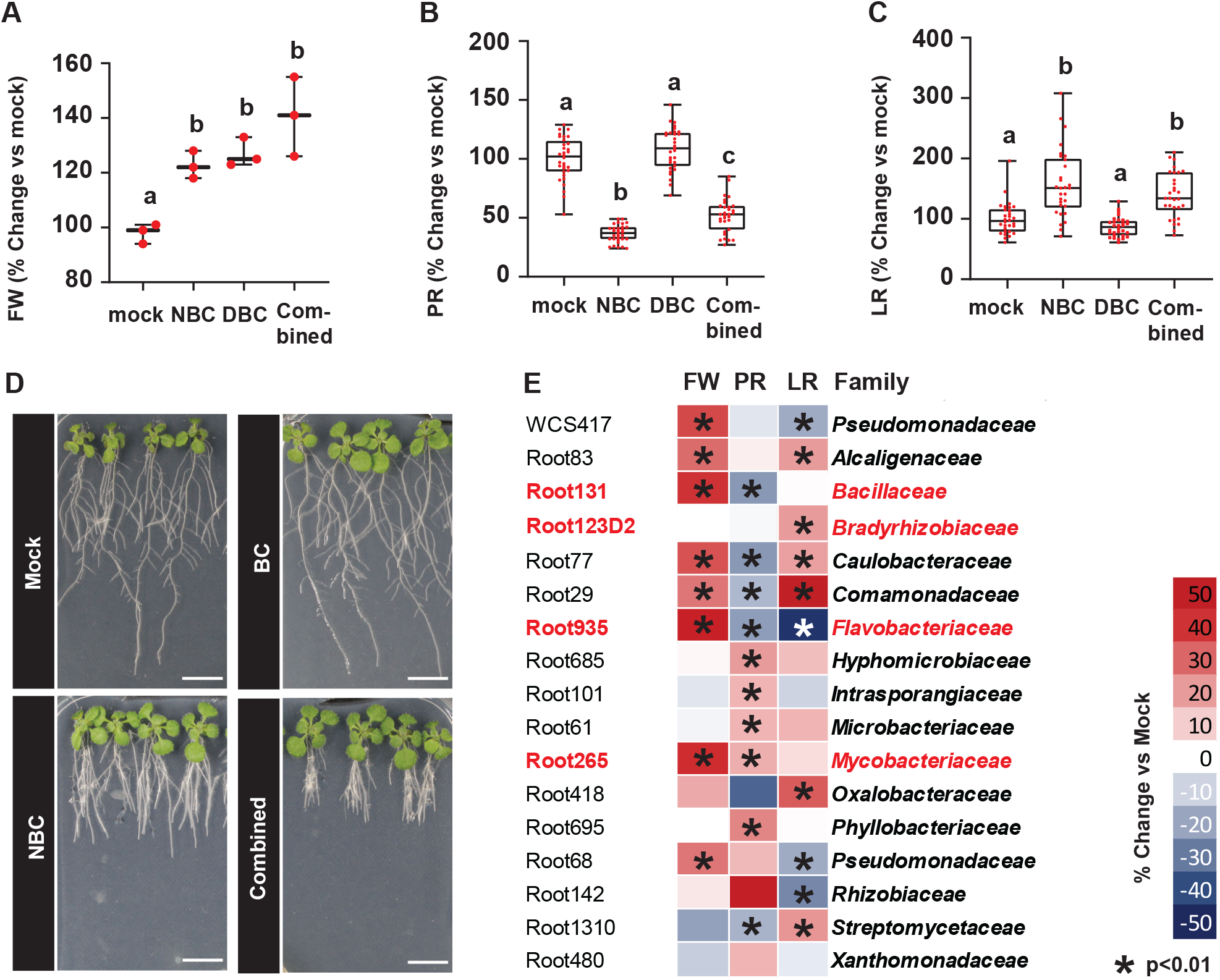
Effects of 16-member synthetic bacterial community and individual strains on plant growth and root system architecture in different plate-assay setups. **A-C)** Effects of a 16-member synthetic bacterial community (SynCom) on Arabidopsis thaliana when grown for 10 days without bacteria (mock), non-bacterial contact with shared headspace (NBC), direct bacterial contact (DBC) or a combination of the two conditions. **A)** Percent changes in total fresh weight normalized to mock average (n=3) (pooled sampling) **B)** Percent changes in primary root length normalized to mock average (n=30) **C)** Percent changes in lateral root occurrence normalized to mock average (n=30). Letters indicate statistically significant differences between groups (one-way ANOVA and Tukey’s post hoc pairwise HSD t-test, p<0.01). **D**) Representative pictures of plants subject to treatments from A-C. Scale bars are 1 cm. **E)** Effects of the Pseudomonas strain WCS417 and individual SynCom members on plant phenotype when grown without direct bacterial contact (NBC). Results depicted as means, including three biological replicates containing 10 seedlings each. The changes are shown in percentage compared to mock control. (*) Asterisks indicate statistically significant differences compared to control via Student t-test (p<0.05). Strains labeled in red were selected for further analysis. FW, fresh weight; PR, primary root length; LR, lateral root occurrence.

### VOCs from individual bacteria have distinct effects on plant organ growth

To disentangle the bVOC effect arising from the SynCom, we measured how individual members of the community affected plants in the NBC configuration. As a positive control, we employed the Pseudomonas strain WCS417, known to trigger growth promotion in Arabidopsis seedlings upon bVOCs exposure (Pieterse et al., 2021; Zamioudis et al., 2015). Plants which shared headspace with the strains WCS417, Root83, Root131, Root77, Root29, Root935, Root265 or Root68 showed a significant increase in the total fresh weight (Figure 1E). Root685, Root101, Root61, Root265, Root695 or Root142 elicited a significant PR elongation, while Root131, Root77, Root29, Root935, Root418, or Root1310 significantly reduced the PR length (Figure 1E). Root83, Root123D2, Root77, Root29, Root418, and Root1310 had a positive effect on lateral root occurrence, whereas WCS417, Root935, Root68 and Root 142 had reduced lateral root numbers when compared to mock-inoculated controls (Figure 1E). Taken together, the individual strains elicited a range of distinct responses affecting the shoot biomass or root architecture that overall differ from that of the SynCom. Thus, the SynCom-specific response may result from dominant key species present in the SynCom, or it may be an emergent property from species influencing each other in a mixed community. To probe this further, we selected Root123D2, Root131, Root935, Root265, which all had a positive or neutral effect on overall plant growth, but triggered differing root responses, for further analysis.

### VOC-mediated shoot growth promotion and root inhibition are strain-dependent

These selected strains were individually tested in NBC, DBC, or combined configurations. Root131, Root935 and Root265, which had positive effects in the NBC configuration (Figure 1E and 2A) also gave rise to increased fresh weight in the DBC and combined configurations, whereas Root123D2 led to increased shoot weight only in DBC configuration (Figure 2A). It is therefore possible that the observed shoot growth promotion by these species occurs through bVOCs, even in conditions where the bacteria are in direct contact with the plant. Interestingly, while the primary root length was decreased in NBC-grown plants grown with Root131 or Root935 (Figure 1E and 2A), Root131 led to increased root length in the DBC configuration and combinatorial treatments (p<0.01, Students T-Test, Figure 2A), whereas Root935 DBC-grown roots were not affected, and the primary root was shorter also in combined treatment. Root935 caused reduced lateral root occurrence in the NBC configuration, while Root265 led to an increased occurence in the DBC configuration (Figure 2A). For all NBC conditions, we observed a dose-dependent effect when inoculated with a series of bacterial dilutions (Supplementary Figure. 2). In summary, bVOCs from the individual members of the growth-promoting SynCom provide distinct effects on root and shoot modulation, which may or may not be prevalent in a community context.

**Figure 2.**
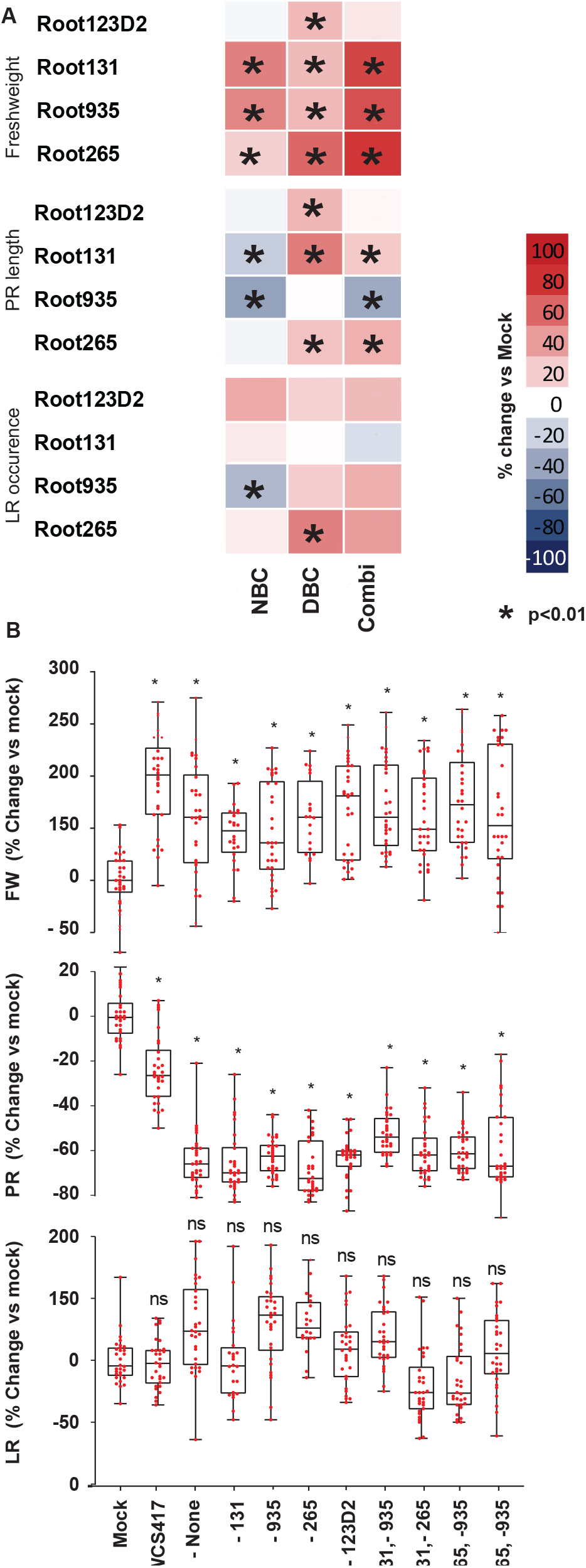
Effects of selected strains and dropout SynComs on plant growth and root system architecture. **A)** Effects of the Pseudomonas WCS417 strain and 4 selected SynCom members on plant growth under 3 different plate setups conditions (Non-Bacteria Contact, with shared headspace (NBC), Bacteria Contact (DBC) and a combination of NBC&D-BC (Combi)) (n=30). The changes are shown in percentage compared to mock control. Results depicted are means of three biologicalreplicates containing 10 seedlingseach. Asterisks (*) indicate a statistically significant difference compared to control (Students t-test, p<0.01) **B)** Relative effects of SynCom drop-out communities on plants grown in non-bacterial conditions with shared headspace. FW, fresh weight; PR, primary root length; LR, lateral root occurrence (n= 28-30). The changes are depicted in percentage comparing each treatment with mock control. Asterisk (*) indicates statistically significant differences compared to mock control. ns, not significant indicates no significant differences compared to mock control.

### Drop-out SynComs highlight possible redundancy in VOC effects on plant growth

To investigate if and how the selected bacterial strains dominantly exert the effects observed on the co-inoculated plants, we performed drop-out experiments, where up to three strains were omitted from the 16-member SynCom. These reduced SynComs were then grown with plants in the NBC configuration. In all drop-out combinations, the fresh weight of exposed plants was significantly higher than the mock-treated control (p<0.01, Students T-Test Figure 2B, top panel). Moreover, we also observed a significant reduction of PR length (p<0.01, Figure 2B, middle panel) whereas no significant effects were seen on lateral root occurrence (Figure 2B, lower panel). It is therefore likely that the effects observed on plants by bVOCs from the full SynCom is an emergent trait of the bacterial community.

### Drop-out SynComs show strain-specific changes in chemical defense responses

As the plants may respond to the presence of bVOCs by inducing defense responses (Weisskopf et al., 2021), we further addressed to what extend the bVOCs affected chemical defense signatures of the plants by measuring changes in accumulation of camalexin and glucosinolates, which are part of the immune response to bacterial and fungal pathogens in the Brassicaceae (Bednarek, 2012). When inoculated individually in the NBC setup, only Root265 induced significant camalexin accumulation in the exposed plants (Figure 3A). However, in plants exposed to the full SynCom, we found that the levels of camalexin were strongly reduced (Figure 3B). This effect was maintained in all drop-out experiments with Root131, Root935, Root265 or Root123D2 removed (Figure 3B). However, and somewhat surprisingly, drop-out combinations that excluded both Root265 and Root935, either a lone ortogetherwith Root131, showed levels of camalexin accumulation similar to that of mock-inoculated controls (Figure 3B). In none of the investigated combinations were aliphatic glucosinolates significantly affected by the bVOCs (Figure 3C and D). However, among the individual strains, bVOCs from Root131 and Root935, as well as WCS417, resulted in decreased accumulation of indolic glucosinolates (Figure 3E). Surprisingly, the full 16-member Syncom did not affect the concentration of indolic glucosinolates, but dropping out any of the strains, alone or in combination, except for Root131, caused a significant decrease in accumulation of this class of glucosinolates (Figure 3F). In summary, we conclude that not only is the SynCom capable of manipulating part of the plant defense system through bVOCs, but intra-community interactions are likely to affect the plant responses. To probe how the overall structure of the community was affected, we characterized the composition of the bacterial communities in the employed drop-out combinations.

**Figure 3.**
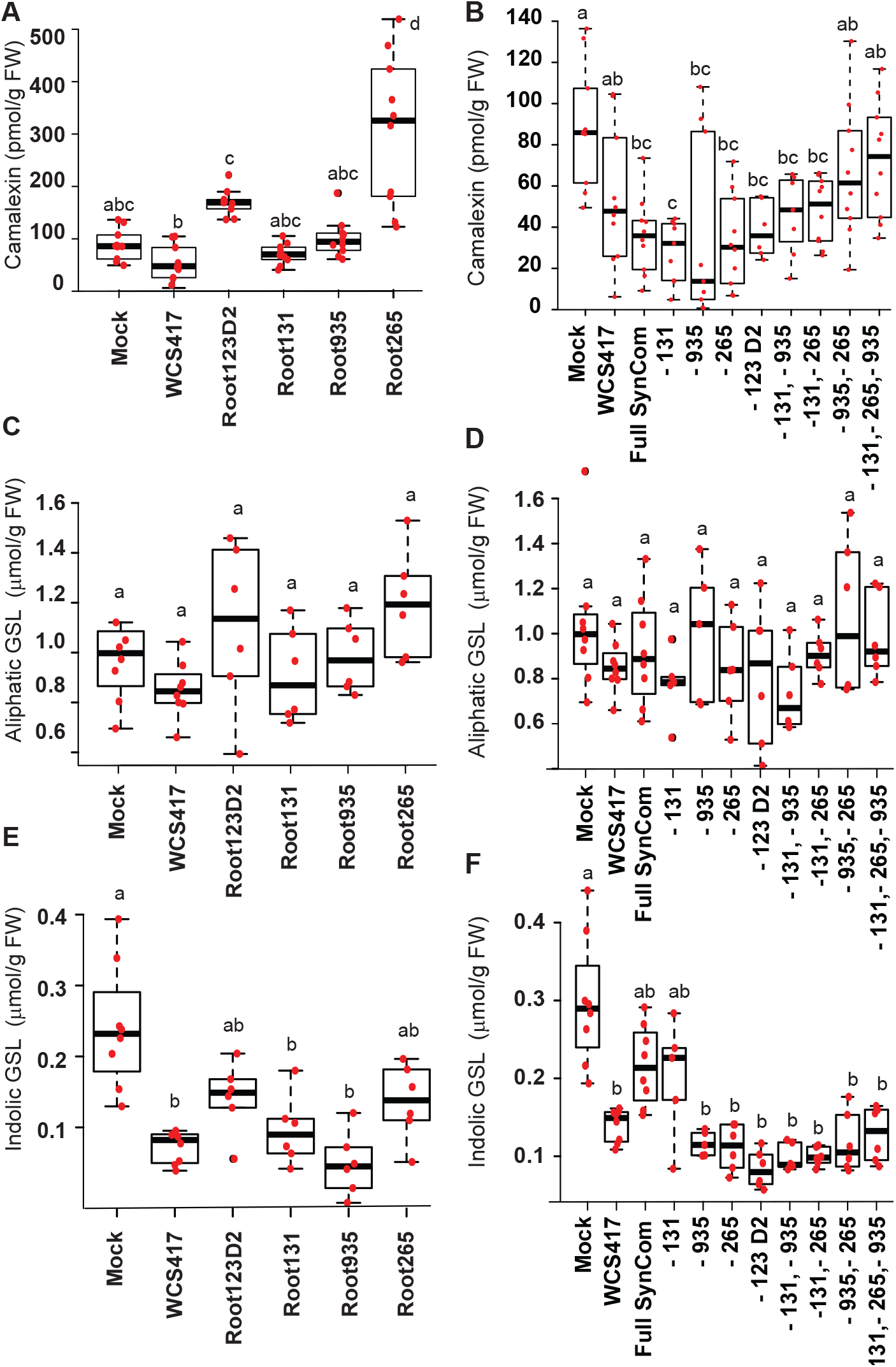
Analysis of plant chemical defense metabolites. Concentration of tryptophan-derived Camalexin (**A and B)**, methionine-derived aliphatic (**C and D)** as well as tryptophan-derived indole glucosinolates (GSL) (**E and F**) in plants grown for 10 days in non-bacterial contact (NBC) configuration with shared headspace. Letters indicate statistically significant differences between the treatment means (one-wayANOVA and Tukey’s HSD Test, p<0.01).

### Community profiles of drop-out SynComs reveal an intricate key-species network

To assess how the removal of individual strains affected the stability of the SynComs grown on agar plates in the NBC condition, we profiled the bacterial community structure via a standardized 16S rRNA amplicon sequencing pipeline (P. Zhang et al., 2021). Principal coordinates analysis of beta-diversity (Bray-Curtis dissimilarities) revealed a separation of SynComs mainly driven by the absence of either Root935 or Root131 (Supplementary Fig. 3A), indicating a major effect of those strains on the overall bacterial community. Calculation of the relative abundance (RA) of individual strains in each SynCom showed patterns that suggested significant changes in the abundance of certain strains in the drop-outs compared to the original SynCom (Supplementary Fig. 2B). However, these may be masked by the relatively high abundance of individual strains (Supplementary Figure 2B). Thus, to quantify this, we removed the sequencing reads for the omitted strain(s) of a given drop-out SynCom from the full-SynCom samples and compared the RA of each strain (Figure 4, Methods). Removal of Flavobacterium Root935 consistently led to a significant (p<0.05) increase of Caulobacter Root77 and Microbacterium Root61. Omitting Mycobacterium Root265 from the communities did not significantly change the RA of any strain. When Bacillus Root131 was removed alone or together with Root265, the Pseudomonas strain Root68 was significantly more abundant, but not when Root935 was also removed. These results indicate that microbe-microbe interactions affect the community structure of these SynComs and highlight this phenomenon as a likely driver of the observed plant responses.

**Figure 4.**
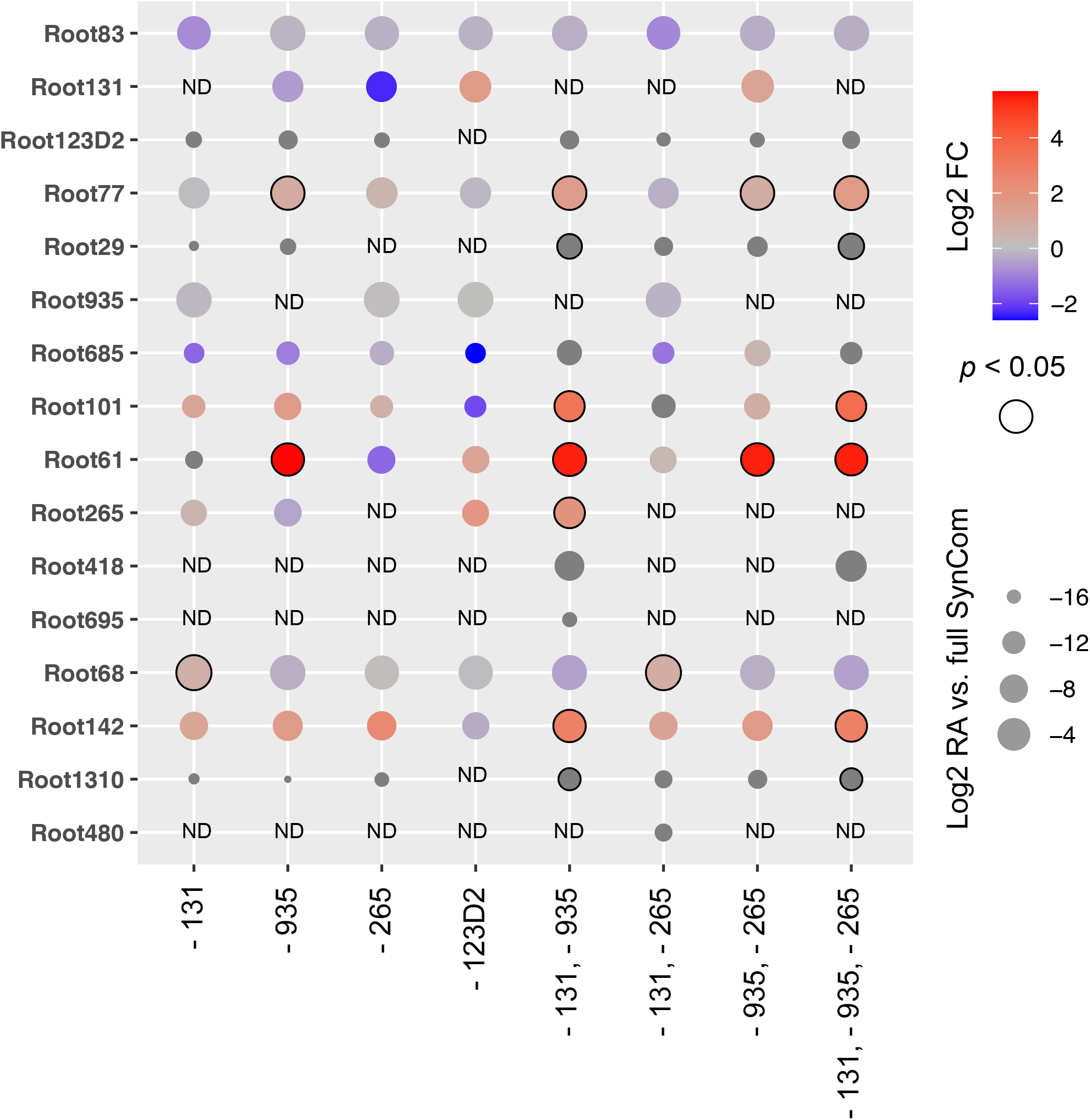
Community profiling of the drop-out SynComs. Specific strains are differentially abundant in drop-out SynComs. Dot plot showing the relative abundance (RA) of each strain in each SynCom condition compared to the full 16-member SynCom. The dot size corresponds to the mean RA. The color indicates the log2 fold change of RA in the drop-out condition relative to the RA in the 16-member SynCom after in silico depletion of the relevant strain(s). A circled dot indicates a significant change (Wilcoxon rank sum test, P<0.05).

## Discussion

Roots sense and integrate signals from the aboveground plant parts and their surrounding environment and use these signals to actively establish host-specific bacterial communities as part of their microbiota (Fitzpatrick et al., 2018; Wippel et al., 2021). The root-associated microbiota can extend the functional repertoire of the host, providing protection against pathogens or facilitating nutrient acquisition, which leads to improved growth (Pascale et al., 2020). The host-specific microbiota structure seems to be crucial for these functions, and a change in composition that negatively affects the plant is generally known as dysbiosis (Paasch & He, 2021). Perturbations of community composition can happen through antibiotic metabolites or photosynthates originating from the host plant (Koprivova et al., 2019) or via antagonistic or cooperative interactions among bacterial species (Jacoby & Kopriva, 2019). While attention has been given to plant metabolites shaping the microbiome, other chemical players in this context, such as the bVOCs emitted by bacterial strains, have been understudied. Indeed, bVOCs from individual strains are able to promote plant growth (C.-M. Ryu et al., 2003; C. M. Ryu et al., 2004), and are therefore likely contributing to these functions. This has important agricultural implication, however, we only have limited insights into how bVOCs influence plant-microbe communication in a bacterial community context.

We set out to investigate the effect of bVOCs released from a growth-promoting SynCom consisting of bacterial strains occurring as part of a natural Arabidopsis root community. Our finding that this synthetic community could induce plant growth without direct bacterial contact with the roots in a dose-dependent manner (Figure 1, Supplementary Figure 1A) suggests that bVOCs may contribute to the plant growth promotion previously reported for this SynCom (Wippel et al., 2021). However, the bVOCs also led to changes in root architectural traits not observed when bacteria were allowed to colonize the roots (Figure 1B and C). Possibly, the plant response to direct contact of bacteria overrides the effects caused by the bVOCs alone. Alternatively, the SynCom grown in a separate media compartment gives rise to different bVOCs than bacteria directly applied to the media of the plant compartment. Indeed, it was previously observed that bVOCs profiles and plant growth modulation depended on the type of bacterial growth medium (Blom et al., 2011). Future studies aiming at determining bacterial physiology under changed media conditions will likely shed light on how these differences in plant performance are mediated.

Even a small-complexity 16-member SynCom offers a very high number of possible plant-bacteria and bacterial-bacteria interactions. This very likely influences the bouquet of bVOCs produced by the community and collaborative or antagonistic interactions in the community may be an imporant part of finetuning bVOC-induced plant responses. Complex communities of bacteria can influence plant performance in a positive or negative manner dependent on presence or absence of distinct bacterial taxa (Finkel et al., 2019; Ramirez-Villacis et al., 2020; Vogel et al., 2021). Moreover, host-associated antagonism and competition play an important role in shaping biotic steady-states (García-Bayona & Comstock, 2018). Thus, we were interested to test, if dominant strains that may drive the bVOC effects observed on the plant growth were being favored under the employed growth conditions. While we did not identify specific community members predominantly responsible for the growth alterations of Arabidopsis (Figure 2B), we were able to identify SynCom members prone to grow only in absence of members known to induce antagonistic behavior (Powers et al., 2015; Simões et al., 2008). In addition, several members were only present in very low amounts or completely undetected after 10 days of growth (Figure 4). Thus, it is likely that antagonistic interactions occur within this natural, albeit strongly simplified community and that this allows modulation of the bVOC profile. In support of this, keystone bacterial species that are needed to secrete certain bVOCs responsible for growth promotion have been identified (Carlström et al., 2019). Metabolic distance showed a significant correlation with antagonism, which was mainly driven by few strains. Thus, it is possible that even small differences in bacterial community composition have an important influence on the overall bVOC profile. Future characterization of bVOCs profiles from the different SynComs will allow to identify links between the bacterial species and a given bVOC bouquet, and the chemical nature of their effect on plant performance.

Modulation of plant immune responses is a prevalent feature of bacterial microbiota members (Hacquard et al., 2017). Indeed, when quantifying the defense-related metabolites camalexin and glucosinolates in plants exposed to bVOCs as a proxy for immune output, it was intriguing to find that this is strongly dependent on the bacterial component (Figure 3A and B). In particular, contactless mono-associations between plants and Root265 led to a strong increase specifically in shoot camalexin production and not indole glucosinolates, whereas this was repressed whenever Root265 was present in a SynCom (Figure 3A, B, E and F). Surprisingly, this repressive effects of the SynCom bVOCs on camalexin induction was lost whenever strains Root265 and Root935 were not included in the SynCom (Figure 3B). Thus, both highly specific immune repressive and inductive effects of bVOCs exist within bacterial communities. Interestingly, Pseudomonadaceae strains (represented in the employed SynCom by Root68) have been shown to actively repress innate immune responses (Millet et al., 2010). Thus, it is possible that this strain may contribute to the repressive effect by bVOCs. A more detailed analysis of plant immune elicitation that include drop-out combinations without Root68 and Root935 may shed light on this. On the other hand, none of the strains was able to induce the accumulation of glucosinolates. This is particularly interesting in the case of Root265, as camalexin and indolic glucosinolates share part of their biosynthetic pathway. Analyses of the effects of bVOCs on expression of genes belonging to this biosynthetic pathway will be helpful to understand the interplay during this chemical defense, and potentially establish a framework to identify the underlying mechanims responsible for this observation.

In summary, our work emphasizes that bVOCs profiles produced by bacterial communities are likely to be an important aspect of plant-microbe communication and shaping bacterial community composition. Our finding that the production of plant chemical defenses is influenced by volatiles from distinct bacterial communities opens up ways for deeper mechanistic understanding of bVOC effects such as defense priming through bacterial presence. Underground effects that influence plant growth and resistance have important agronomical implications. The ability to manipulate plant performance via microbe interactions, and in particular bVOCs, represents an exciting novel opportunity to improve plant health and may lead to a deeper molecular understanding of the mechanisms that drive overall plant performance in the field.

## Acknowledgments

TGA thanks the Max Planck Society (MPG) and the Sofja Kovalevskaja programme of the Alexander von Humboldt foundation for funding. KW is funded through the DFG (German Research Foundation) priority programme SPP 2125 DECRyPT. SK and AK research is funded by the DFG under Germany’s Excellence Strategy – EXC 2048/1 – project 390686111 and SPP2125 DECRyPT – project 401836049. We thank Irene Klinkhammer for technical assistance with metabolite analyses, Paul Schulze-Lefert and the “Rooters” for insightful comments on the work.

## Author contributions

GMT designed and conducted all growth experiments and analyzed the data, RC initiated the project and designed growth experiments, AK and SK designed, conducted and performed chemical analysis, KW designed, conducted and analyzed bacterial profiling experiments. TGA designed and supervised experiments and wrote the manuscript with KW and GMT. All authors read and approved the manuscript.

## Data avaliabilty

All datasets are avaliable from the corresponding authors upon request.

---

**Supplementary Figure 1:**
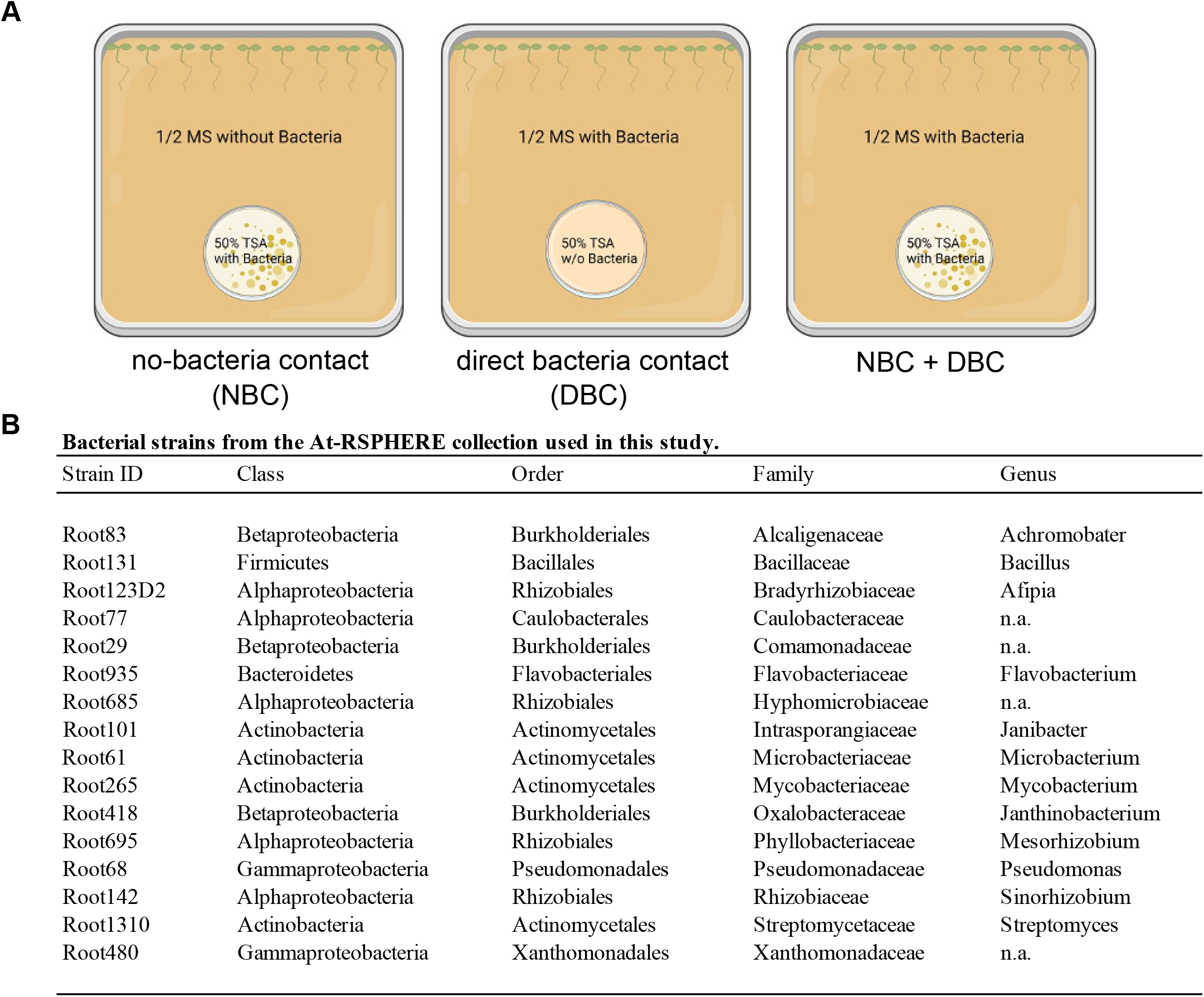
Experimental setup and strains used in this study. A) Schematic showing the plant and bacterial growth configurations. Created with Biorender.com B) List of bacterial strains used in this study. Strains were originally isolated from roots of A. thaliana grown in natural soil.

**Supplementary Figure 2:**
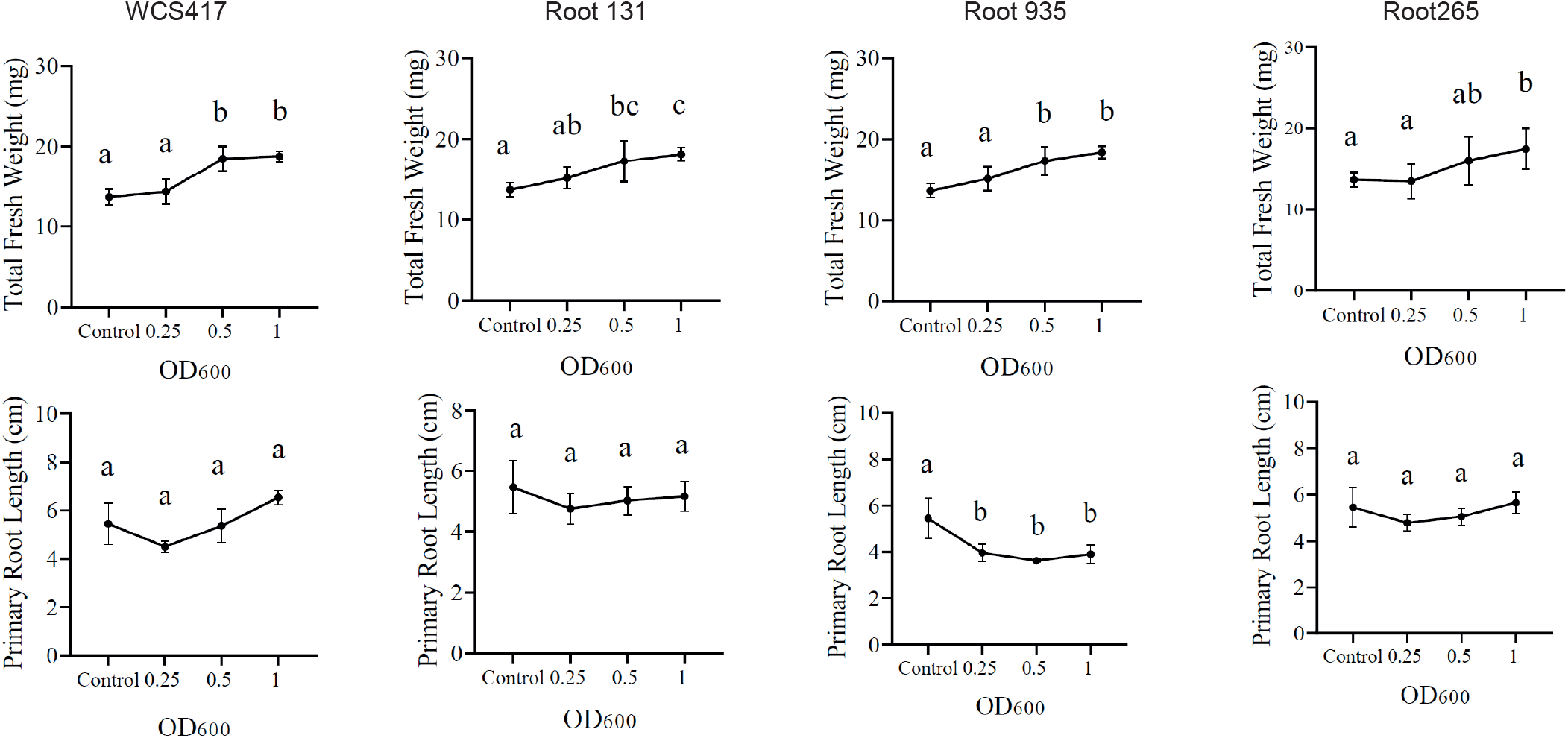
Effects of bacterial density on plant performance in the NBC configuration. Effects of different bacterial densities on A. thaliana total fresh weight (mg) and primary root length (cm) inoculated with bacterial strains using an OD600 of 0.25, 0.5, and 1. Results represent the means of 3 biological replicates, each containing 10 plants.Letters indicate a statistically significant difference between the treatment means (one-way ANOVA and Tukey’s HSD Test, p<0.05).

**Supplementary Figure 3:**
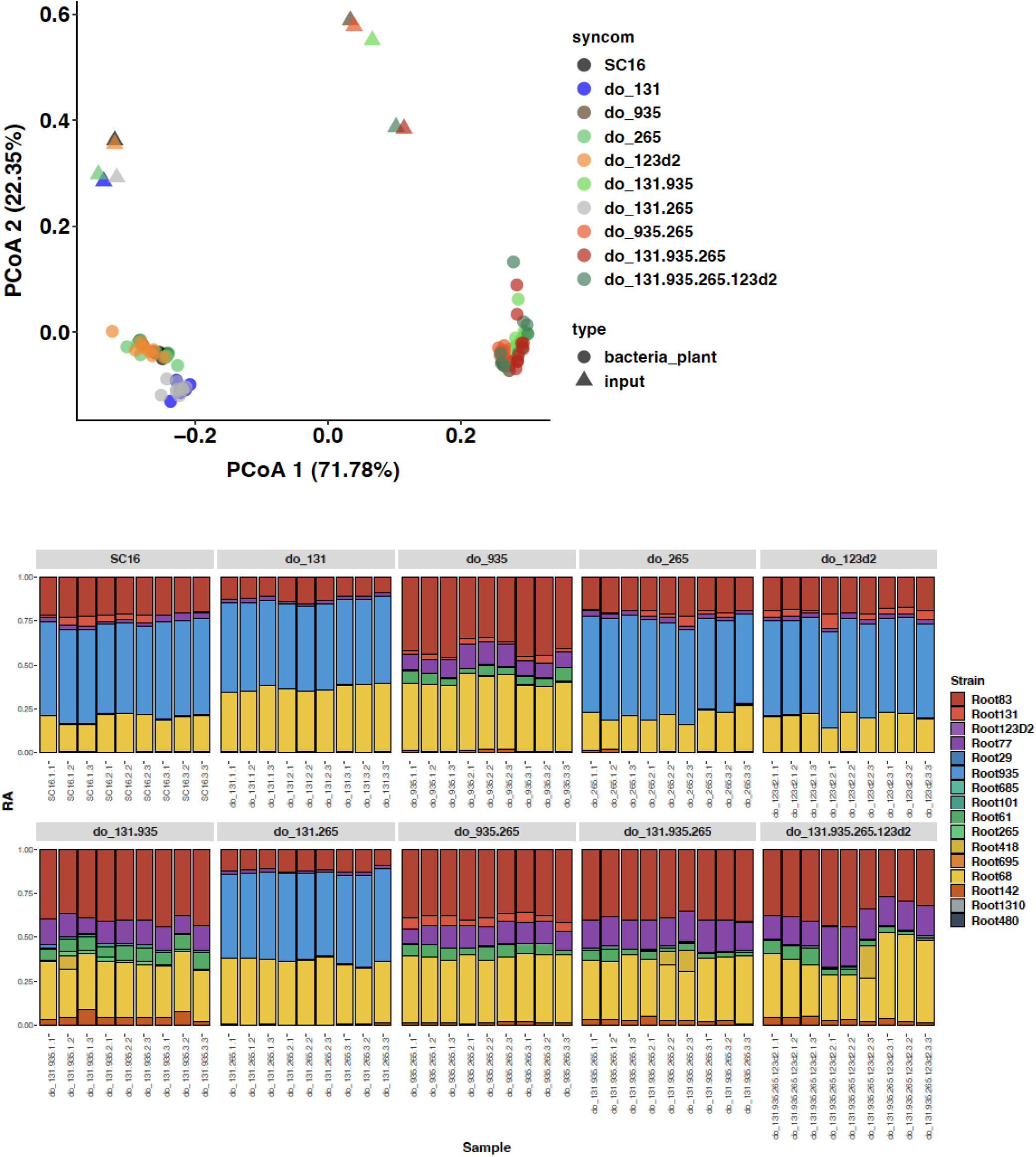
Diversity of bacterial drop-out SynComs. A) Principal coordinates analysis of Bray-Curtis dissimilarities of bacterial SynComs grown on agar plates in a split-plate system for non-contact growth of plants and bacteria. Input samples are the SynCom preparations at the start of the experiment, and output communities are the bacterial samples after 10 days of growth next to the plants. SynComs are either the 16-member community (SC16), or the drop-out (do_) communities with omitted strains indicated by their ID number. B) Bar plots showing the relative abundance (RA) of individual strains in the indicated SynComs in each replicate.

